# Systems therapeutics analyses identify genomic signatures defining responsiveness to allopurinol and combination therapy for lung cancer

**DOI:** 10.1101/396697

**Authors:** Iman Tavassoly, Yuan Hu, Shan Zhao, Chiara Mariottini, Aislyn Boran, Yibang Chen, Lisa Li, Rosa E. Tolentino, Gomathi Jayaraman, Joseph Goldfarb, James Gallo, Ravi Iyengar

## Abstract

The ability to predict responsiveness to drugs in individual patients is limited. We hypothesized that integrating molecular information from databases would yield predictions that could be experimentally tested to develop genomic signatures for sensitivity or resistance to specific drugs. We analyzed TCGA data for lung adenocarcinoma (LUAD) patients and identified a subset where xanthine dehydrogenase expression correlated with decreased survival. We tested allopurinol, a FDA approved drug that inhibits xanthine dehydrogenase on a library of human Non Small Cell Lung Cancer (NSCLC) cell lines from CCLE and identified sensitive and resistant cell lines. We utilized the gene expression profiles of these cell lines to identify six-gene signatures for allopurinol sensitive and resistant cell lines. Network building and analyses identified JAK2 as an additional target in allopurinol-resistant lines. Treatment of resistant cell lines with allopurinol and CEP-33779 (a JAK2 inhibitor) resulted in cell death. The effectiveness of allopurinol alone or allopurinol and CEP-33779 were verified in vivo using tumor formation in NCR-nude mice. We utilized the six-gene signatures to predict five additional allopurinol-sensitive NSCLC lines, and four allopurinol-resistant lines susceptible to combination therapy. We found that drug treatment of all cell lines yielded responses as predicted by the genomic signatures. We searched the library of patient derived NSCLC tumors from Jackson Laboratory to identify tumors that would be predicted to be sensitive or resistant to allopurinol treatment. Both patient derived tumors predicted to be allopurinol sensitive showed the predicted sensitivity, and the predicted resistant tumors were sensitive to combination therapy. These data indicate that we can use integrated molecular information from cancer databases to predict drug responsiveness in individual patients and thus enable precision medicine.

## Introduction

Lung cancers are the most common cause of death related to cancers worldwide (*1*). Non-small cell lung cancer (NSCLC) is a widely occurring lung cancer that includes three main subtypes: adenocarcinoma, squamous cell carcinoma and large cell carcinoma (*2*). Targeted treatment for lung cancer based on attacking the major mutational characteristics and responsiveness to immunotherapy has significantly increased life span (*3-5*). Often, many of the mutated gene products that are drivers of the cancers are part of, and controlled by, complex networks of cellular components within cancer cells. Such cellular regulatory networks give rise to the biological capabilities that are characteristic of cancer cells (*6*). In spite of the steady advances in the treatment of lung cancers, a targeted therapy often works only on a subset of patients with the target driver mutation. One approach to search for other possible therapies, rests with the possibility that many pathways are uniquely dysregulated in individual patients, and these pathways can be used to find targets for potential efficacious drugs. Systems level analyses that consider different types of omics data can provide both the breadth and depth needed to identify pathways that can be targeted therapeutically. Such analyses can also enable the discovery of prognostic genomic biomarker sets associated with the therapeutic targets, and thus represent an important step in precision medicine.

We used a combination of cancer databases for data integration to identify specific drugs that are effective in a predictable manner in individuals. We started with The Cancer Genome Atlas (TCGA) (*7*) to test our hypothesis that integrated consideration of the molecular characteristics of individual patient tumors will allow us to identify actionable drug targets. TCGA contains both clinical and molecular data from individual patients for different types of cancers including lung cancer. These data have led to reclassification of many cancers based on molecular characteristics (*8-10*). We focused on lung adenocarcinoma (LUAD). We explored TCGA data from LUAD patients to find new pathways and targets which had not previously been used for drug therapy of lung cancer. Our strategy was to focus on targets that are not well-known mutations or that have protein kinase activity, to be able to explore unidentified potential cancer genes(*11*). We found the xanthine dehydrogenase (*XDH)* gene highly expressed in a subset of patients with lower survival rates in TCGA LUAD data. *XDH* and its interconvertible form xanthine oxidase have been known as drug targets for over fifty years. In fact, an inhibitor of XDH, allopurinol, was synthesized and tested as an early potential anti-cancer agent. Although allopurinol was a not a successful anti-leukemic drug (*12*), it has been used successfully to treat gout for over fifty years and is also used to prevent kidney stones associated with hyperuricemia caused by cancer chemotherapy (*13, 14*). We then experimentally analyzed LUAD cell lines in the Cancer Cell Line Encyclopedia (CCLE) (*15*) to identify cell lines that are either sensitive or resistant to allopurinol. We used molecular data associated with these cell lines to identify the transcriptomic signatures that predict sensitivity or resistance to allopurinol. We also used network analysis to predict that cell lines resistant to allopurinol alone could be successfully treated with combination therapy of allopurinol with a JAK2 inhibitor. We tested this prediction experimentally and found it to be valid. We then used the molecular signatures from integration of TCGA and CCLE data to analyze transcriptomic data from patient-derived tumors (PDX) from Jackson laboratory and were able to identify tumors that had allopurinol sensitivity indicating that molecular signatures for allopurinol sensitivity can be identified.

## Results

### Analysis of TCGA and identification of XDH as a target in NSCLC

We analyzed methylation and gene expression data (12905 genes) from patients with lung adenocarcinoma (LUAD) in TCGA. All differentially expressed genes and methylation markers were identified. The genes which were common to both the set of differentially expressed genes and the set of differentially expressed methylation markers were selected (25 genes). These 25 genes then were evaluated for correlation with “days to death” for individual patients. Among these 25 genes, we found 16 genes which had either positive correlations between “days to death” and DNA methylation signatures or negative correlations between “days to death” and gene expression signatures. Of these 16 genes, 4 of them were selected as novel and druggable: *AGMAT* (Agmatinase), *ATIC* (5-Aminoimidazole-4-Carboxamide Ribonucleotide Formyltransferase/IMP Cyclohydrolase), *FAM83A* (Family With Sequence Similarity 83 Member A) and *XDH* (Xanthine Dehydrogenase). We selected *AGMAT*, *ATIC* and *XDH* for experimental validation because all had enzymatic activity while *FAM83A* has no known enzymatic function. (Figure 1-A and Figure 1-S1)

**Figure 1:**
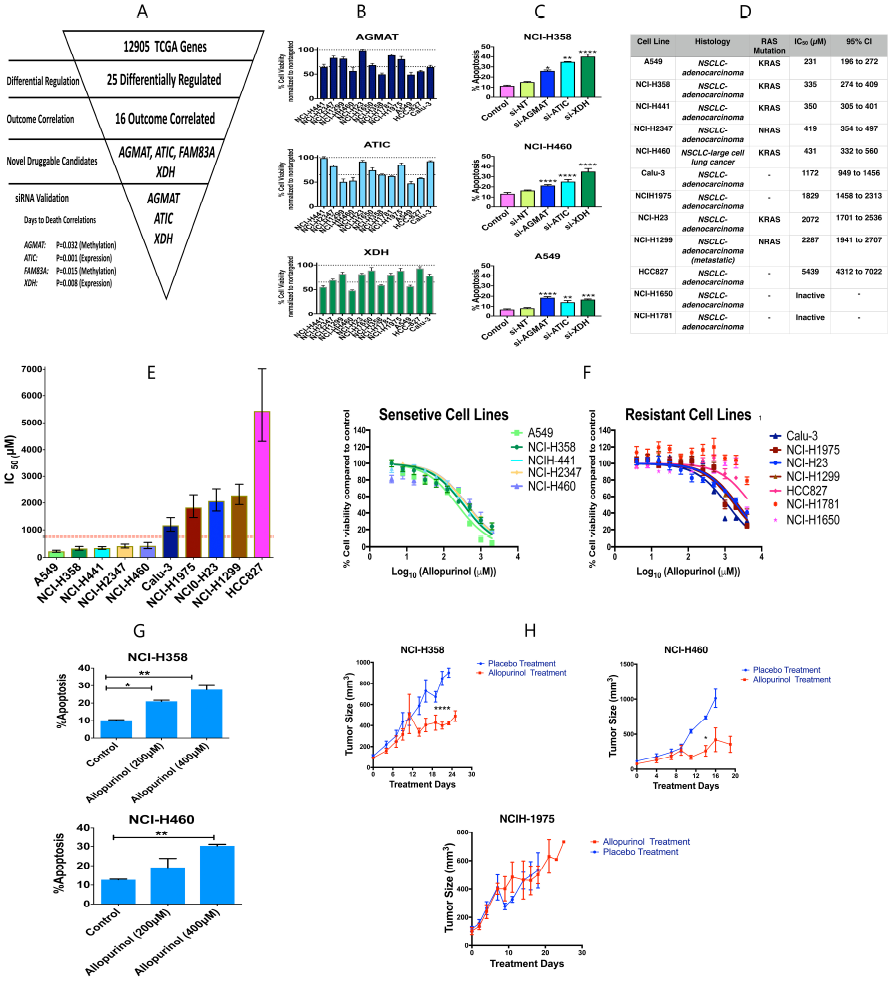
A) Analysis of 12905 genes in TCGA led to finding 25 differentially regulated genes in patients with lung adenocarcinoma, of these genes, 16 correlated with clinical outcome. Four of these 16 genes were novel druggable genes and 3 of them were selected for siRNA knockdown validation. B) siRNA knockdown validation of 3 gene targets in 12 NSCLC cell lines, the results are shown as percent cell viability (mean±SEM). C) Percent of apoptosis in cells (expressed as percent of Annexin V positive cells) induced by siRNA knock-down of these three gene targets in 3 NSCLC cell lines (mean+SEM). siRNA knock-down increased apoptosis compared to control. (One-way ANOVA, *p<0.05, **p<0.01, ***p<0.001, ****p<0.0001). D) List of 12 cell lines, their histology, their RAS mutation status, the allopurinol IC_50_ for reduction of viability and its 95% confidence interval E) Comparing allopurinol IC_50_s in the different cell lines (error bars are 95% confidence intervals). An IC_50_ < 754μM (red dotted line) was chosen as the criterion for considering a cell sensitive to allopurinol. F) Log concentration-response plots for cell lines sensitive to and resistant to allopurinol. (mean±SEM) G) Allopurinol-induced apoptosis (expressed as percent of Annexin V positive cells) in NCI-H358 and NCI-H460 cell lines in a concentration-dependent manner (mean+SEM, One-way ANOVA, *p<0.05, **p<0.01). H) Xenograft models were used to assess the effect of allopurinol on 2 cell lines sensitive to allopurinol (NCI-H358 and NCI-H460) and one cell line resistant to allopurinol (NCI-H1975). Allopurinol (200mg/kg) was administered by oral gavage 3 times a week to treatment groups (n=15 for NCI-H358, n=10 for NCI-H460 and n=8 for NCI-H1975) and PBS by oral gavage to the placebo groups (n=7 for NCI-H358, n=4 for NCI-H460 and n=9 for NCIH1975). The tumor size in mice bearing NCI-358 and NCI-H460 cells and receiving allopurinol was significantly decreased compared to placebo at day 23 and 14 respectively. (mean±SEM, unpaired t-test, *p<0.05, ****p<0.0001) Allopurinol did not have any significant effect on tumor size of mice bearing NCI-H1975 cells.

Using a panel of twelve NSCLC cell lines from CCLE, we evaluated the effect on cell viability of knocking down the expression of *AGMAT*, *ATIC* and *XDH* genes by siRNA. Immunoblotting analyses showed that all siRNAs efficiently suppressed expression of each of the genes tested (Figure 1-S2-A) indicating that the screen results were “on-target”. The siRNA knock-down experiments showed differing cell viability in cell lines subjected to gene knock-down (Figure 1-B). To further investigate the role of these genes in cell survival, we used siRNA gene knockdown on 3 of the cell lines (NCI-H358, NCI-H460 and A549) each of which had its viability decreased by a third or more after knockdown of each of the 3 genes. Apoptosis, indicated by percent of Annexin V positive cells, was significantly induced by knock-down of each of these 3 genes compared to non-targeting siRNA (Figure 1-C). Knock-down of each of these 3 genes also resulted in changes in cell cycle phases. Knock-down of *XDH* in NCI-H358 and NCI-H460 cells significantly increased the cells arrested in G2/M phase compared to control and non-targeting siRNA (Figure 1-S2-B)

### Allopurinol-sensitive and allopurinol-resistant phenotypes in NSCLC

We selected *XDH* for subsequent investigations, because there is already an FDA-approved drug, allopurinol, that inhibits the enzyme, and that failed as an anti-neoplastic drug when it was first synthesized and tested. (*12*). Based on our correlation analysis of TCGA data, *XDH* was among those genes whose higher level of expression correlated with lower survival rates for a subset of patients. Hence, we reasoned that inhibiting *XDH* could potentially change the course of cancer cell progression. We treated our panel of twelve NSCLC cell lines with allopurinol and calculated the IC_50_ of allopurinol for cell viability. Figure 1-D shows list of these cell lines, their histology, IC_50_ of allopurinol and its 95% confidence interval and the presence of RAS mutations. All of these cell lines were adenocarcinoma. Rank ordering of mean IC_50_s revealed a greater than 2 fold increase between NCI-H460 (431 μM) and Calu-3 (1172 μM), and their 95% confidence intervals did not overlap. We thus chose an IC_50_ =754μM (midway between the upper bound of the confidence interval for NCI-H460 and the lower bound of the confidence interval for Calu-3 as the dividing line between sensitivity and resistance to allopurinol (Figure 1-D and E). Of the five sensitive cell lines, 4 were positive for KRAS mutations, and one for NRAS. The log-concentration-response plots for the 5 sensitive cell lines, and the 7 resistant cell lines are presented in Figure 1-F. Allopurinol could also, in a concentration-dependent manner, induce apoptosis in sensitive cell lines (Figures 1-G and Figure 1-S3-A) and it also could increase the percentage of cells arrested in G2/M phase of the cell cycle compared to vehicle-control (Figure 1-S3-B).

Basal level of expression of XDH protein in these 12 cell lines negatively correlated with IC_50_ for allopurinol (Spearman r=-0.8667, P=0.002) which implies an addiction to XDH protein in sensitive cell lines (Figure 1-S4).

We then tested two allopurinol sensitive (NCI-H358 and NCI-H460) and one allopurinol-resistant cell line (NCI-H1975) in an NCR-nude mouse xenograft model. Administration of allopurinol (200mg/kg by oral gavage, three times a week) for 30 days reduced the tumor size significantly in mice bearing NCI-358 and NCI-H460 cells at day 23 and 14 compared to placebo group, but had no significant effect on tumor size in mice bearing NCI-H1975 cells (Figure 1-H) (Comparisons were made on the last day all mice in each group were alive; some of the mice either died or were sacrificed prior to 30 days based on IACUC protocols for treatment of animals). Expression of cleaved caspase 3 in tumor samples after 30-day allopurinol treatment showed apoptosis induction in sensitive cells engrafted in mice compared to placebo groups. There was no remarkable increase in cleaved caspase 3 levels in tumors formed by NCI-1975 cells, which are allopurinol-resistant (Figure 1-S5-A). Immunofluorescence images of tumor samples from xenograft models revealed allopurinol-induced apoptosis indicated by cleaved caspase 3 expression as well as decreased blood vessel density indicated by CD31 expression (Figure 1-S5-B). Figure 1-S6 shows that there was no significant changes in the body weight for the xenograft models of all treatment regimens.

### Genomic signatures of responsiveness to allopurinol

A key mutation differentiating allopurinol-sensitive and allopurinol-resistant cell lines appears to be KRAS. However, it is neither necessary nor sufficient. KRAS mutations were found in 4 out of 5 sensitive cell lines (the remaining one had an NRAS mutation), but in only 1 out of 7 resistant cell lines (one other of which had an NRAS mutation). Given this variability, we reasoned that additional molecular determinants could provide a more predictive signature. To find the genomic determinants of responses of these cell lines to allopurinol, we used CCLE transcriptomic data to find differentially expressed genes in allopurinol-sensitive and allopurinol-resistant cells. We used a t-test (p < 0.001) to identify the genes with the highest expression levels in each group (Figure 2-A and B and Figure 2-S1). Gene set enrichment analysis using these sets of signatures revealed possible pathways involved in responses to allopurinol treatment (Figure 2-S2). In sensitive cells, pathways and processes related to oxidative stress were most prominent. Reactive oxygen species (ROS) and their metabolic functions are possible pathways regulating this response. Previous integrative analysis of TCGA data of NSCLC (*7*) has shown that alterations of oxidative stress pathways are among the recurrent aberrations of key regulatory processes in lung adenocarcinoma. In resistant cells, fatty acid catabolic processes and other functions related to metabolism of lipids are likely to be dominant. Gene signatures (combined and separately) also were subjected to pathway enrichment analysis using Molecular Biology of the Cell Ontology (MBCO) (*16*), (Figure 2-S2-A). Bar diagrams visualize the negative log_10_(p-values) (Fisher’s Exact Test) of the top 5 predicted sub-cellular processes (SCPs) of the levels 1 (brown), 2 (red) and 3 (blue) (*16*). Among processes enriched for combined signatures (all 12 genes) were cellular responses to stress, lipid metabolism, cellular responses to oxidative stress, and the JAK-STAT signaling pathway.

**Figure 2:**
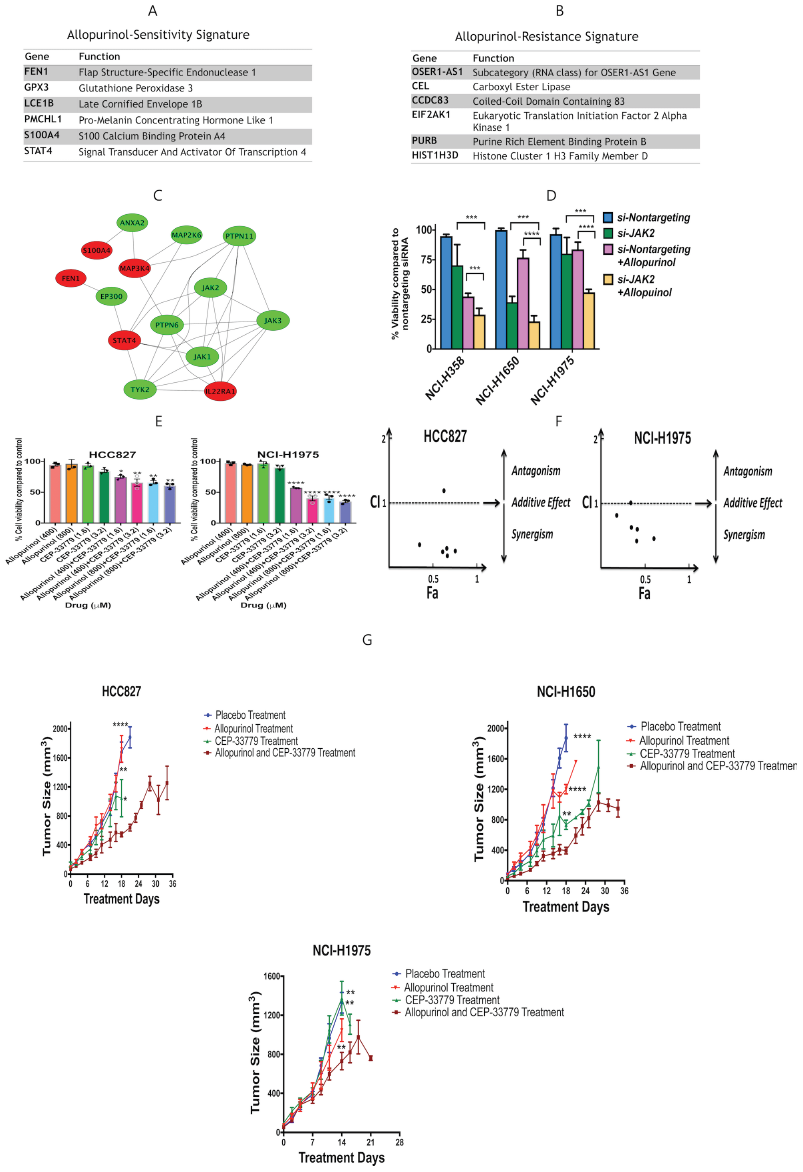
A and B) Genomic signatures of sensitivity and resistance to allopurinol. Each genomic signature includes a set of 6 genes considered to have high expression in either sensitive or resistant cell lines. The sensitive phenotype is also characterized by the presence of a RAS mutation.C) Protein interaction network built using genomic signatures. Relaxation of stringency led to a gene set capable of forming a network with additional intermediary nodes like JAK2. D) Validation of combination treatment with allopurinol (400μM) and knockdown of *JAK2* in 3 cell lines, percent of cell viability compared to control is shown. Combination treatment significantly decreased cell viability compared to allopurinol treatment alone and *JAK2 knockdown alone*. (mean+SEM, One-way ANNOVA test, ***p<0.001, ****p<0.0001) E) Effects of combination treatment with CEP-33779 and allopurinol on cell viability of NCIH1975 and HCC827 cell lines. Combination therapy significantly decreased cell viability compared to allopurinol treatment alone. Percent of cell viability compared to vehicle-control is shown. (mean±SEM, one-way ANNOVA, *p<0.05, **p<0.01, ****p<0.0001) F) Combination Index (CI) for different doses of CEP-3379 and allopurinol in NCI-H1975 and HCC827 cell lines. CI lower than 1 indicates a synergistic effect; which is the case for most of the combination doses. G) Combination therapy with CEP-33779 and allopurinol in xenograft models using 3 different cell lines. PBS, allopurinol (200mg/kg) alone, CEP-33779 (10mg/kg) alone and combination doses of allopurinol (200mg/kg) and CEP-33779 (10mg/kg) were administered by oral gavage 3 times a week. [placebo group (n= 6 for HCC827, n=6 for NCI-H1650 and n=5 for NCI-H1975), allopurinol treatment group (n= 6 for HCC827, n=6 for NCI-H1650 and n=5 for NCI-H1975), CEP-33779 treatment group (n=7 for HCC827, n=6 for NCI-H1650 and n=6 for NCI-H1975) and combination therapy group (n=7 for HCC827, n=7 for NCI-H1650 and n=8 for NCI-H1975)]. The tumor size in mice bearing HCC827 and NCI-H1650 and receiving combination therapy was significantly decreased compared to placebo and single treatment groups at day 18. The tumor size in mice bearing NCI-H1975 and receiving combination therapy was significantly decreased compared to placebo and single treatment groups at day 14 (mean+SEM, unpaired t-test, *p<0.05, **p<0.01, ****p<0.0001)

Based on the pathways enriched by gene signatures and the known biochemistry of XDH activity in purine metabolism and redox balance in cells (*12*) (Figure 2-S3-A), we built a toy computational model of addiction to XDH. The observation of higher XDH protein expression in allopurinol-sensitive cell lines (Figure 1-S4) suggests that the XDH level regulates reprogramming of metabolic dependency of lung adenocarcinoma cells. Figure 2-S3-B presents a toy mathematical model to explain this phenomenon using arbitrary parameters and a fuzzy membership function (*17*). Increased XDH protein expression (Figure 1-S4) is assigned to dependency on the pentose phosphate pathway (PPP) which balances higher reactive oxygen species (ROS) levels produced by XDH activity. This balance is important for cell survival, as otherwise, increased ROS will induce apoptosis. Inhibiting XDH should dramatically affect PPP and this perturbation leads to cell death because cells are addicted to PPP. This is the case in allopurinol-sensitive cells, however, resistant cells have lower levels of XDH which makes them not require PPP because they can use fatty acid catabolism instead. Reducing XDH activity cannot cause cell death in these cells.

### Combination therapy: Allopurinol with a JAK2 inhibitor

We used network analysis to find additional drug targets in allopurinol-resistant cells. Statistical cutoffs with high stringency (p=0.001) resulted in small lists of genes (6 genes in each category) that do not form networks. Relaxation of stringency and selecting the top 10 down-regulated and the top 10-up-regulated genes from differentially expressed genes led to a larger list (20 genes) that could form networks with the addition of intermediary nodes from the human protein-protein interaction network (Figure 2-C). One of these nodes was JAK2 which is a target for FDA approved drugs. For example, ruxolitinib is approved for treatment of myelofibrosis and tofacitinib is approved for treatment of rheumatoid arthritis and psoriatic arthritis (*18-20*). JAK2 inhibitors like baricitinib, gandotinib and lestaurtinib are being tested in clinical trials for a variety of diseases including acute myeloid leukemia (*21-23*). Another line of evidence for the role of JAK2 in responsiveness to allopurinol is the gene set enrichment analysis of gene signatures (Figure 2-S2). We significantly decreased JAK2 protein expression (Figure 2-S4-A) using siRNA knock-down in one allopurinol sensitive cell line (NCI-H358) and two allopurinol-resistant cell lines (NCI-H1650 and NCI-H1975). Allopurinol treatment (400μM) after *JAK2* gene knock-down significantly decreased cell viability compared to treatment with allopurinol alone or *JAK2* gene knock-down alone, indicating boosting and restoration of the sensitive phenotype (Figure 2-D). Treatment of resistant cells (HCC827 and NCI-H1975) with a combination of allopurinol and a JAK2 inhibitor (CEP-33779) significantly decreases cell viability compared to treatment with allopurinol alone. Treatment with a combination of allopurinol and CEP-33779 is synergistic. We calculated the combination index (CI) (*24*) for 6 different combinatory concentrations of allopurinol and CEP −33779 and the resulting CI was less than 1 in 5 combinations in each of these 2 cell lines indicating a synergistic effect (Figure 2-F). CI analysis also shows synergism of allopurinol and CEP-33779 in three other allopurinol-resistant cell lines (Figure 2-S5). These synergistic effects were observable even at low concentrations of CEP-33779 (1.6 and 3.2μM) combined with allopurinol (Figure 2-E and F and Figure 2-S5). Combination treatment also significantly diminished colony formation for both an allopurinol‐ sensitive (NCI-H358) and an allopurinol-resistant (NCI-H1975) cell line in soft agar gel compared to vehicle-control (Figure 2-S6). These observations indicate that combination therapy with allopurinol and CEP-33779 also can boost the anti-neoplastic effect of allopurinol in sensitive cells.

We evaluated the effect of combination therapy with allopurinol and CEP-33779 on xenograft models with three resistant cell lines (Figure 2-G and 2-S7). NCR-nude mice bearing HCC827, NCI-H1650 and NCIH-1975, cells were administered placebo (phosphate buffered saline (PBS)), allopurinol (200mg/kg), CEP-33779 (10mg/kg), or the combination 3 times per week by oral gavage. The tumor size in mice bearing HCC827 and NCI-H1650 and receiving combination therapy was significantly decreased compared to placebo and single treatment groups at day 18. The tumor size in mice bearing NCI-H1975 and receiving combination therapy was significantly decreased compared to placebo and single treatment groups at day 14 (Comparisons were made on the last day all mice in each group were alive; some of the mice either died or were sacrificed prior to 36 days based on IACUC protocols for treatment of animals)). (Figure 2-G and 2-S7)

### Gene signatures are capable of predicting responsiveness to allopurinol and combination therapy

The gene signatures derived from CCLE data were used to evaluate their predictive capability on additional NSCLC cell lines in the CCLE. For this we used a scoring system which was able to rank CCLE cell lines based on gene signatures and assign a quantitative characteristic to them for defining likelihood of sensitivity to allopurinol (Figure 3-A). This algorithm was used to identify the most likely resistant and sensitive NSCLC cell lines in CCLE. These cell lines are listed in Figure 3-B and C. Using a cell viability assay, we calculated the IC_50_ of allopurinol in each of these cell lines. Among the 5 cell lines predicted as sensitive, three had a RAS mutation, and all had an allopurinol IC_50_ less than 754μM. None of the four predicted resistant cell lines had any RAS mutation, and all had an allopurinol IC_50_ greater than 754μM. (Figure 3-B and C) The cell line with the highest IC_50_ for allopurinol in the sensitive group, HCC15 was a squamous cell lung carcinoma, all others were adenocarcinomas, the cell type used to establish the IC_50_ cutoff. Figure 3-D shows log concentration-response curves in both sensitive and resistant cell lines. Except for NCI-H2106, the three other predicted allopurinol-resistant cell lines showed a synergistic effect of combination treatment with allopurinol and CEP-33779. Figure 3-E presents the comparison of cell viability after treatment with allopurinol and CEP-33779 alone and 4 combinatory doses of both drugs in COR-L105 and NCI-H1568 cell lines. Combination of both drugs significantly decreased cell viability compared to single treatment with allopurinol. Figure 3-F shows the CI for 6 combinatory doses of these drugs in these 2 cell lines indicating synergism, and Figure 3-S1 shows the CI for the NCI-H2170 cell line.

**Figure 3:**
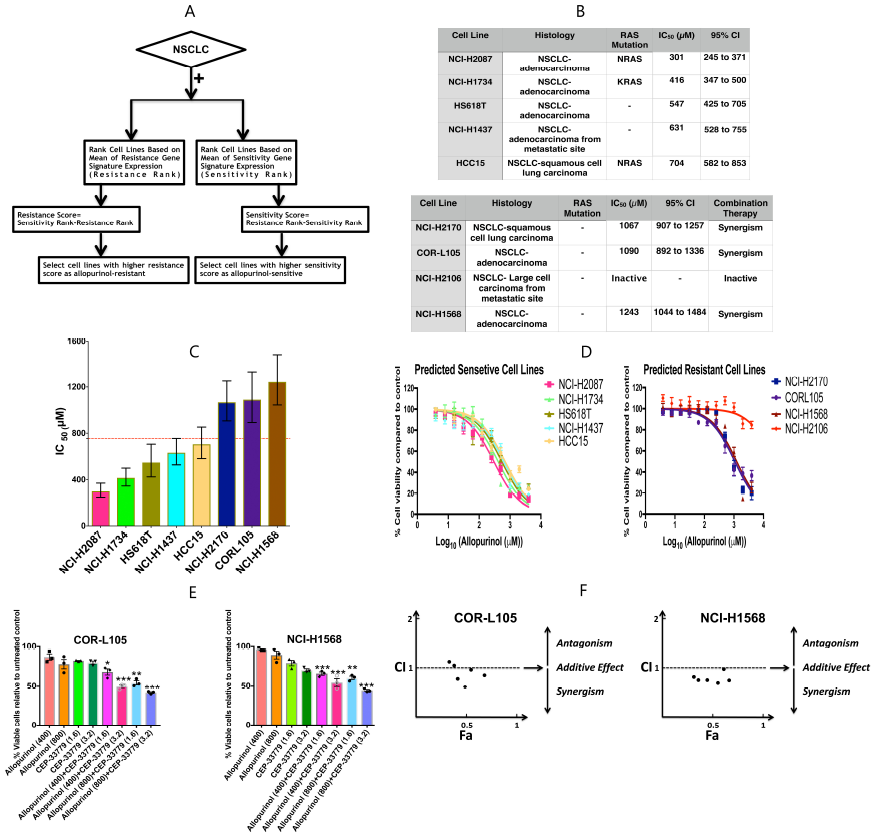
A) Flowchart describing the process of selecting new cell lines as allopurinol-sensitive and allopurinol-resistant. B) List of predicted cell lines, their histology, RAS mutation status, calculated IC_50_ for allopurinol and its 95% confidence interval. C) IC_50_ comparison for the predicted cell lines shown as mean±SEM; error bars are 95% confidence intervals. The IC_50_ cut-off to determine sensitive and resistant cells was considered to be 755μM (red dotted line). D) Concentration-response curves for allopurinol treatment in predicted sensitive and resistant cell lines (mean±SEM). E) Effects of combination treatment with allopurinol and CEP-33779 on two cell lines predicted as resistant (COR-L105 and NCIH1568). Combination therapy significantly decreased cell viability compared to treatments with allopurinol alone. Percent of cell viability compared to vehicle-control is shown. (mean+SEM, one-way ANNOVA, *p<0.05, **p<0.01) F) CI for different concentrations of allopurinol and CEP-33779 in COR-L105 and NCI-H1568 cell lines; most of the combinations showed synergistic effects (CI < 1).

### Gene signatures and the allopurinol sensitivity in PDX models of NSCLC

To test the efficacy of allopurinol and combination therapy with CEP-33779 in patient-derived xenograft (PDX) models of NSCLC, and to evaluate the power of gene signatures to assign tumors for best treatment options, we used the gene signatures of sensitivity and resistance to allopurinol in the PDX models of NSCLC provided by The Jackson Laboratory. We analyzed the gene signatures in two sets of data available from the Jackson Laboratory NSCLC PDX models: RNAseq data was used to select one PDX model as allopurinol sensitive and Affymetrix gene expression data was used to select one PDX model as allopurinol-sensitive and one as allopurinol-resistant. We used an algorithm similar to cell line selection by defining a score for sensitivity and resistance based on gene signatures, and we also considered the RAS mutation status (Figure 4-A). In this algorithm, carrying a KRAS or NRAS mutation was a boolean function to select sensitive models, as RAS mutation was previously observed in most sensitive cell lines. These PDX models were grafted as subcutaneous tumors in NSG mice, and allopurinol (70 mg/kg daily), CEP-3379 (10 mg/kg daily) and combination therapy (Allopurinol 50 mg/kg and CEP-33779 2.5 mg/kg daily) were administered by oral gavage for 30 days. Control mice received placebo (PBS) daily. In models predicted to be allopurinol-sensitive (TM01563 selected using RNAseq data and TM00206 selected using Affymetrix data), treatment with allopurinol alone, significantly decreased the tumor size in mice bearing them compared to the placebo-treated group. After 30 days of treatment, tumor weights in the allopurinol group were significantly lower than those in the placebo group (Figure 4-B and C).

**Figure 4:**
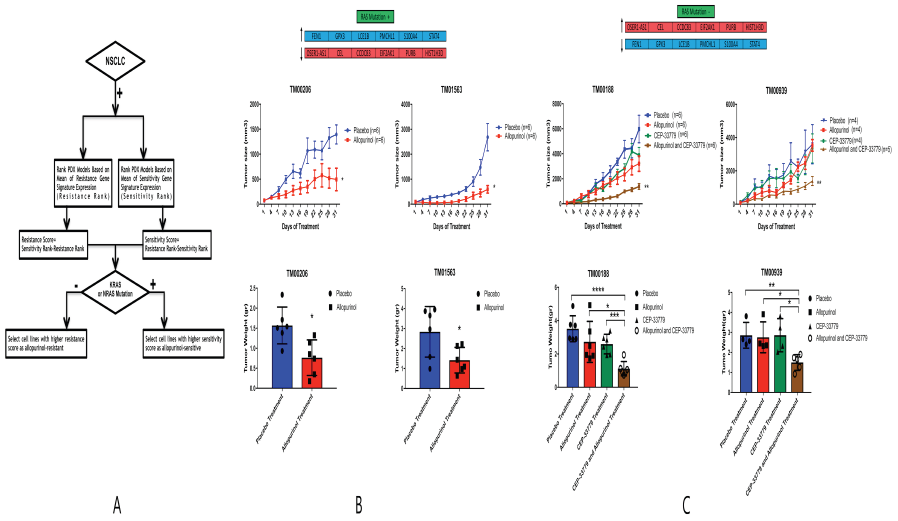
A) Flowchart describing how the sensitive and resistant PDX models were selected from a set of available PDX models. B, C and D) Post-treatment tumor size and tumor weight in 4 PDX models; TM01563 and TM00206 models were predicted as sensitive to allopurinol and TM00188 and TM00939 models were predicted as resistant to allopurinol based on the genomic signatures. Tumor size was significantly lower in the allopurinol group compared to placebo group and at the end of the study tumor weights were significantly lower in the allopurinol group compared to placebo group in TM01563 and TM00206 models. Combination therapy decreased the post-treatment tumor size and tumor weight significantly in the TM00188 and TM00939 models compared to single treatment with allopurinol while allopurinol alone and CEP_33779 alone were not able to decrease the tumor size compared to placebo (PBS). (mean+SEM, unpaired t-test, *p<0.05, **p<0.01, ***p<0.001) (Allopurinol (70 mg/kg daily), CEP-3379 (10 mg/kg daily), combination therapy (Allopurinol 50 mg/kg and CEP-33779 2.5 mg/kg daily) and PBS as placebo daily)

Mice bearing TM00188 and TM00939 model tumors (predicted as allopurinol-resistant using Affymetrix data and RNAseq data respectively), showed a significant decrease in tumor size after receiving combination therapy compared to the placebo group while neither allopurinol alone nor CEP-33770 alone had a significant effect on tumor size compared to the placebo group. After 30 days of treatment, tumor weights in the combination therapy group were significantly lower than in placebo, allopurinol alone and CEP-33779 alone groups (Figure 4-D). Figure 4-S1 shows the images of 3 tumors from each treatment group after sacrificing the mice at the end of treatment, showing the size reduction in allopurinol groups for allopurinol-sensitive tumors (TM01563 and TM00206) and combination therapy groups for allopurinol-resistant tumors (TM00188 and TM00939).

Cleaved caspase 3 was expressed in TM01563 (sensitive) tumors receiving allopurinol and in TM00188 (resistant) tumors receiving combination therapy indicating apoptotic cell death (Figure 4-S2).

Using a pan-caspase *in vivo* assay revealed activation of caspases as indicators of cell death in mice bearing allopurinol-sensitive tumors and receiving allopurinol. This assay also showed that mice bearing allopurinol-resistant tumors receiving combination therapy had more activation of caspases and more cell death (Figure 4-S3). Drug treatment did not lead to any significant weight loss of weights in mice bearing tumors (Figure 4-S4).

Taken together, these data indicate that genomic signatures derived from TCGA can correctly predict allopurinol and allopurinol/CEP-33779 responsiveness in patient derived tumors.

## Discussion

TCGA has been very useful in developing molecular classifications of cancer subtypes that underlie key concepts of precision medicine. In addition, as TCGA contains both molecular and clinical data, it is possible to analyze the relationship between these classes of data to develop predictive signatures for progression of cancers in individuals. Additionally, as our study shows, TCGA data sets are a gold mine for identifying targets for new drugs, drug repurposing and combination therapy. A fundamental premise for such mining is that a particular drug therapy is likely to be effective in only a subset of patients. The transcriptomic signature provides a clear way to identify these patients. While our study was being completed, several papers have described the potential value of transcriptomic signatures. Shukla et al. (*25*) have published a computational study using TCGA data to identify a four gene transcriptomic signature that predicts survival in the TCGA cohort. Li et al. (*26*) have used a combination of transcriptomic data sets to identify an individualized immune signature for prediction of survival. These two studies have focused on predicting survival without specifying the nature of the drug therapy. Lee et al. (*27*) used transcriptomic signatures to predict repositioning of drugs for cancer therapy. Although superficially similar, our approach differs from these studies in the following important ways. In our initial search of TCGA we considered all molecular changes including genomic and epigenomic variations individually, not just transcriptomic changes, for predictions (Figure 1-A).

We subsequently focused on gene expression levels, as this was a facile way to integrate TCGA and CCLE data, using the signatures to identify individual NSCLCs, both in the CCLE cell lines and the PDX tumors. We used network building and analyses to identify relationships between drug targets. The two approaches we have used should be broadly useful in identifying additional drug targets and predicting responsiveness for these drugs in other cancers as well. Combining identification of targets for a drug with specification of which patients might benefit from treatment with that drug, can be a substantive step forward in precision medicine

We focused on allopurinol both for historical and practical reasons. As described by Elion in her Nobel Prize essay (*28*) allopurinol was among the earliest potential anti-cancer drugs synthesized in the 1950’s. Although some of the original biochemical reasoning from over fifty years ago for focusing on allopurinol remains valid today, anchoring its use in molecular characteristics of an individual’s tumor enables accurate prediction of drug sensitivity. Thus, it appears that systems biology approaches have enabled the rediscovery of allopurinol as an anticancer drug. At a practical level, allopurinol is a relatively safe and inexpensive FDA approved drug which could readily be tested in the clinic for patients whose molecular profiling indicates that it could be effective.

Our data indicate that two characteristics predict effectiveness: a RAS mutation and a precision transcriptomic signature are both necessary for treatment of NSCLCs with allopurinol and for combination therapy with allopurinol and a JAK2 inhibitor The role of RAS mutation in NSCLC has been shown to be related to heterogeneity in metabolic dependencies and metabolic reprogramming (*29, 30*). It has been shown that transcriptomic profiles of tumors present a metabolic heterogeneity among individual patients which needs to be taken into account for designing precision therapeutics (*31*). In our cell line studies, most. but not all, sensitive cell lines had a RAS mutation and most, but not all, resistant cells lacked a RAS mutation. This fact indicates the need for a dual signature set including both RAS mutation and gene expression pattern for assignment for treatment with allopurinol or allopurinol plus a JAK2 inhibitor. Although the histology of all cells and tumors in this study was NSCLC, the TCGA data were from adenocarcinoma tumors and all but one of the cell lines used to find the signatures were also adenocarcinoma while some of the predicted cell lines and PDX models were not adenocarcinoma. Interestingly, one of the cell lines predicted as sensitive with squamous cell carcinoma histology was at the border of sensitivity and resistance compared to other predicted cell lines which were adenocarcinoma.

This study starts with data from individual patients and ends with predictive treatment of tumors from individual patients. Although cost considerations prevented us from testing a large number of PDX tumors, the two tumors we predicted would be allopurinol sensitive were shown to be so. This combination of computational predictions and experimental testing demonstrates the potential power of integrating molecular data in both TCGA and CCLE when used in training sets to predict responsiveness of new patients. We anticipate that clinical decision support systems that integrate molecular characteristics with clinical outcomes can become a useful tool for drug selection for individual patients and an important component of a precision medicine strategy in cancer.

## Methods

### Ethics Statement

All animal experiments adhered to a protocol approved by the Institutional Animal Care and Use Committee (IACUC) at the Icahn School of Medicine at Mount Sinai and were performed according to the Office of Laboratory Animal Welfare (OLAW, National Institute of Health) and Animal Welfare Act (AWA, United States Department of agriculture) guidelines.

### Materials

*AGMAT, ATIC, XDH*, *JAK2* and scrambled siRNA were purchased from GE Healthcare., Inc. Anti-XDH antibody was purchased from Sigma-Aldrich. Anti-GAPDH, Cleaved-Caspase 3, and JAK2 were purchased from Cell Signaling Technology, Inc. Anti-CD31 antibody was purchased from Fisher Scientific Company. Allopurinol was purchased from Cayman Chemical Company and CEP-33779 was purchased from Selleck Chemicals LLC. For *in vitro* experiments, allopurinol and CEP-33779 were dissolved in Dimethyl Sulfoxide (DMSO) and then diluted in complete medium to a final DMSO concentration less than 1%. For *in vivo* experiments, allopurinol and CEP-33779 were diluted in phosphate buffered saline (PBS).

### Cell Lines

The human NSCLC cell lines HCC827, NCI-H1437, NCI-H1734, NCI-H358, NCI-H1781, NCIH2170, NIC-H1650, NCI-H2160, NCI-H2087, NCI-H2347, NCI-H441, Hs 618.T, NCI-H1299, NCI-H460, NCI-H1975, NCI-H1568, NCI-H23, Calu-3 and A549 were obtained from the American Type Culture Collection (ATCC, Manassas, VA). The human NSCLC cell line CORL105 was purchased from Sigma-Aldrich and human NSCLC cell line HCC-15 was purchased from Creative Dynamics. The HS618.T cell line was cultured in Dulbecco’s Modified Eagle’s medium (DMEM), supplemented with 10% fetal bovine serum (FBS). A549 was cultured in F-12K medium, supplemented with 10% FBS. NCI-H2160 was cultured in HITES medium supplemented with 5% FBS. Calu-3 was cultured in Eagle’s Minimum Essential Medium (EMEM) supplemented with 10% FBS. NCI-H2087 was cultured in RMPI-1640 medium supplemented with 5% FBS. All other cell lines were maintained in RMPI-1640 medium supplemented with 10% FBS. Cells were grown at 37°C in a humidified 5% CO_2_:95% air atmosphere.

### Cell Cycle and Cell Viability Assay and Calculation of IC_50_ and CI

For cell cycle analysis, cells were fixed in 70% ethanol. Fixed cells were treated with RNase for 20 minutes before addition of 5μg/mL Propidium Iodide (PI) and analyzed by FACS. Cell viability was detected by luminescent cell viability dye (CellTiterGlo, Promega Corporation, USA). Cells were seeded in triplicate into 96-well plates in full growth media. After 24 hours, drugs of interest (allopurinol and/or CEP-33779) were added in 12 different concentrations (varying from 0 to 4mM) and after 48 hours of drug treatment, 20μL of dye was added to each well containing 100μL of treated media. Cell viability was calculated by dividing each luminescent reading by the average of the luminescent readings obtained for vehicle-control. Concentration-response curves were generated and fitted in Prism 7.0 (GraphPad Software, Inc., USA). The IC_50_ values were generated using the log inhibitor-normalized response variable slope function: Y=100/(1+10^((X-LogIC_50_))). IC_50_ values are shown with 95% confidence interval from at least three independent experiments. To evaluate synergism, CI values were calculated based on the method proposed by Chou and Talalay (*24*) using CompuSyn software (*32*). The following single doses of allopurinol were used: 400μM, 800μM and 1000μM. The following single doses of CEP-33779 were used: 1.6μM, 3.2μM and 16μM. The following combination doses were used: Allopurinol=400μM combined with CEP-33779 (1.6μM, 3.2μM and 16μM) and Allopurinol=800μM combined with CEP-33779 (1.6μM and 3.2μM and 16μM).

### Xenograft Cell Line *in vivo* Experiment

NCR-nude female athymic mice were purchased from Taconic Farms, Inc. Mice were injected in the flank region with 1.5*10^6^ cells, while anesthetized with a combination of ketamine and xylazine. Size of tumors was measured in three dimensions using a caliper and tumor volume was calculated by this formula: V=0.5*length*width*height. When tumors reached a minimum size of 100mm^3^, mice were randomly assigned to treatment groups and drug treatment was administered by oral gavage. Allopurinol (200mg/kg three times a week) and CEP-33779 (10 mg/kg three times a week) were diluted in PBS for treatment groups and PBS was given to the control group as placebo. Tumors and weights of the mice were measured 3 times a week.

### Colony Formation in Soft Agar

Cells (1×10^5^ to 2×10^5^ per plate) were suspended in soft agar containing 5% serum, dosed with vehicle and drugs and allowed to grow for 2 to 3 weeks with periodic dosing to keep the dosing media fresh and the agar hydrated. Viable colonies were stained with Iodonitrotetrazolium Chloride at 0.5 mg/mL overnight. Colonies larger than 0.3 mm in each field were manually scored using a light microscope.

### Immunofluorescent and Western Blot analysis of Tumor Tissue

Mice bearing subcutaneous tumors were sacrificed after the treatment course and tumors were resected. These resected tumors were snap-frozen in isopentane, submerged in liquid nitrogen and sectioned onto positive slides. Unstained frozen sections were fixed for 15 minutes in ice-cold acetone, dried, rehydrated in PBS and blocked in Tris-buffered saline (TBS) containing 1% Bovine Serum Albumin (BSA), 10% goat serum followed by overnight (4°C) incubation with primary antibodies for caspase 3 and CD31. After washing, Alexafluor 568-Goat anti-rabbit secondary antibodies (Fisher Scientific Company) were incubated with the tissue for 1 hour at room temperature, followed by 4’,6-diamidino-2-phenylindole (DAPI) (Molecular Probes) staining. Staining was visualized using an Olympus MVX10 Macroview microscope with a 2X Apochromat lens with 5× zoom. For western blot analysis, a 2 to 3 mm cross-sectional slice of the tumor was lysed in RIPA buffer by sonication and the resulting lysates were analyzed by western blot following standard methods.

### PDX Models *in vivo* Experiments

PDX models were purchased from the Jackson Laboratory and they were received as a single tumor engrafted subcutaneously in a NSG mouse (The NOD.Cg-Prkdc^scid^ Il2rg^tm1Wjl^/SzJ). This original mouse was sacrificed and the tumor was divided and engrafted in 5 other NSG mice subcutaneously and allowed to grow. Then each of the new tumors was engrafted in 5-10 more mice. Drug treatments were started at passage 4 when enough tumor samples were available. All NSG mice were purchased from the Jackson Laboratory. When tumor sizes were between 50-150 mm^3^, mice bearing tumors were randomly assigned to treatment groups. Each group had at least 8 mice at the beginning of the experiments. Drug preparation, administration and tumor measurements were the same as in the xenograft cell line *in vivo* experiment, but the allopurinol, CEP-33779, and combination therapy were applied at the following doses: allopurinol (70 mg/kg daily), CEP-33779 (10mg/kg daily) and combination therapy (allopurinol 50mg/kg daily+CEP-33779 2.5mg/kg daily). Tumors and weights of the mice were measured 3 times a week. After the treatment course (30 days), 3 mice from each group were used for *in vivo* imaging using Pan Caspase (VAD-FMK) near infrared assay (Vergent Bioscience) in The IVIS^®^ Spectrum *in vivo* imaging system.

### Statistical Analysis

All experimental data are shown as mean±SEM. Unpaired t-test and one-way ANOVA were used and p<0.05 was considered as significance. All statistical analyses of experimental data were done in GraphPad Software 7.

### TCGA Candidate Gene Signature Identification

In brief, we identified patients with both methylation data and gene expression data from the TCGA-LUAD dataset (Snapshot 12/2012). In addition, we excluded any patient who did not have tissue level control samples. We divided samples into four categories: case-male, case-female, control-male, and control-female. We evaluated the significance of the difference between case-control by determining the absolute difference among the mean divided by the square root of the sum of variance among each of the groups for each gene. This is given by the formula below:

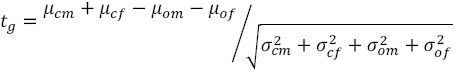

In this formula *t_g_* is the significance of a gene expression, μ_x_ is the average gene expression for the specified category, σ_x_ is the standard deviation of the gene expression for the specified category. The gene expressions were determined by the Agilent 4502A microarray. We selected for gene targets that had a positive gene expression change where *t_g_* > 1.5.

We repeated this procedure selecting for methylation markers assessed by Illumina Human Methylation 27k microarray. In this procedure we selected for markers that had a *t_g_* < −2. We repeated the procedure for markers assessed by Illumina Human Methylation 450k microarray. We identified methylation patterns that were selected based on both array formats. We then selected genes that were selected by both gene expression and methylation differentials.

Upon identifying gene signatures of interest, we correlated the gene expression signatures and methylation signatures to the “days to death”. We used linear correlations to evaluate the associations between molecular data and clinical data. A gene signature was identified as correlated if it had a correlation coefficient that had a one-sided *P* < 0.05 as evaluated by the student’s T distribution. Through this, we identified genes that had either positive correlations between “days to death” and DNA methylation signatures or negative correlations between “days to death” and gene expression signatures. (Supplemental Files 1, 2, and 3)

### Extracting the Gene Signatures of Sensitivity and Resistance to Allopurinol

CCLE gene expression data of 12 cell lines tested for siRNA screening were used. A Welch’s ttest, with P<0.001 was used to compare the differentially expressed genes in two sets of cell lines of allopurinol sensitive and allopurinol-resistant. 12 genes were found (6 in each set) which were differentially expressed.

### Network analysis of Gene Signatures

To find a gene set capable of forming a protein-interaction network, we selected the top 10 upregulated genes and top 10 down-regulated genes. We used X2K to build a protein interaction network using these new gene sets.

### Selecting Cell Lines for Validation of Gene Signatures

For predicting new cell lines as allopurinol-sensitive and allopurinol-resistant, we extracted CCLE gene expression data of all NSCLC cell lines. We then used mean of normalized expression of all genes in gene signature of allopurinol sensitivity to rank all of these cell lines; this rank of cell lines was called sensitivity rank. We also used mean of normalized expression of all genes in gene signature of allopurinol resistance to rank all of these cell lines, this rank of cell lines was called resistance rank. We calculated sensitivity score as Sensitive Score=Resistance Rank-Sensitive Rank and we calculated resistance rank as Resistance Score=Sensitivity Rank-Resistance Rank. We selected the top 5 cell lines (those available to purchase) with highest sensitivity score as allopurinol-sensitive cell lines (if a cell line was not available, we used the next available cell line in the ranked list of allopurinol-sensitive cell lines). The same method was used to select allopurinol-resistant cell lines and 4 cell lines were selected for validation *in vitro*.

### Selecting PDX Models for Validation of Gene Signatures

We extracted gene expression and RAS mutation data of all NSCLC PDX models provided by the Jackson Laboratory. We analyzed the data based on the technology used for measuring gene expression separately (Supplemental files 4, 5 and 6). There were 35 NSCLC PDX models available with RNAseq expression data and 18 NSCLC PDX models available with Affymetrix hg10st gene expression data. For selecting allopurinol-sensitive models, we first calculated the sensitive score and resistance score the same way we calculated them for the cell lines. Then among models with highest sensitivity score and positive for RAS mutation, we selected a model for validation as an allopurinol-sensitive model using RNAseq and Affymetrix hg10st gene expression data (TM01563 model and TM00206 Model respectively).

Among models with highest resistance score and negative for RAS mutation, we selected a model for validation as an allopurinol-resistant model using Affymetrix hg10st gene expression data (TM00188).

### Gene Set Enrichment Analysis

The sets of gene signatures (allopurinol-sensitivity and allopurinol-resistance) were used for gene set enrichment analysis by Enricher (Go ontology and WikiPathways) and MBC ontology(*16, 33*).

### Fuzzy Metabolic Switch Model

A form of the Wilson-Cowan equation (*34*) was used as a fuzzy member function to generate a toy model of a fuzzy metabolic switch as follows:

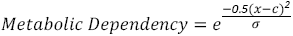

For the toy model presented in figure 2-S3, x was considered to be XDH protein level, σ=1.5, 1 and 0.9 and c=0.1, 0.05 and 2.9. The model was simulated in MATLAB R2017a.

## Acknowledgements

This research was supported by the Systems Biology Center grant GM-071558 and by GM54508. SZ was supported by the Integrated Training in Pharmacological grant GM062754

---

Figure 1-S1

The pipeline used to analyze TCGA data combining molecular alterations and clinical outcome to find new targets which are determined by the clinical outcomes in patients.

Figure 1-S2

A) Western blots showing the protein levels of 3 selected gene targets after siRNA knockdown in 2 of the 12 cell lines tested.

B) Comparison of cell cycle phases in 2 cell lines after knockdown of *AGMT*, *ATIC* and *XDH* compared to control. *XDH knock*-down increased cells arrested in G2/M phase (one-way ANOVA, *p<0.05, **p<0.01, ***p<0.001, ****p<0.0001).

Figure 1-S3

A) Apoptosis induction by allopurinol shown by detection of cleaved caspase 3 in NCI-H358 and NCI-H460 cell lines. B) Compared to control-vehicle, allopurinol arrested the cells (NCI-H358 and NCI-H460) in G2/M phase shown by cell cycle analysis using flow cytometry.

Figure 1-S4

A) Basal XDH protein levels in the NSCLC cell lines. B) XDH protein levels negatively correlate with the IC_50_ for allopurinol in cell lines (Spearman r=-0.8667, P=0.0022). Cell lines sensitive to allopurinol have higher levels of XDH protein indicating an addiction to XDH protein.

Figure 1-S5

A) Allopurinol-induced apoptosis presented as expression of cleaved caspase 3 in xenograft models of allopurinol-sensitive cell lines (NCI-H358 and NCI-H460) but not in NCI-1975 which is allopurinol-resistant. T1-T3 show three different tumor samples. B) Immunofluorescence images of xenografts from PBS and allopurinol-treated mice. Apoptosis induced by allopurinol is indicated by cleaved caspase 3 expression while decreased blood vessel density is indicated by CD31expression.

Figure 1-S6 Changes in body weight for the xenograft models of all treatment regimens.

Figure 2-S1 Number of genes in genomic signatures (genomic signature size) is determined by stringency of statistical analysis on genomic profiles of samples (cell lines). By increasing the p value, the genomic signature size increases. Using p<0.001 leads to a genomic signature with size of 12 (sensitivity signature size of 6 and resistance signature size of 6). Panel A shows changes of size of genomic signature (Resistance and Sensitivity together) for different p values. Panel B shows changes of size of sensitivity genomic signature (red line) and resistance genomic signature (blue line) for different p values.

Figure 2-S2 Gene set enrichment analysis of genomic signatures of allopurinol sensitivity and resistance using GO terms (A and B, for sensitivity genes set and resistance genes set respectively) MBC ontology (C) and WikiPathways (D).

Figure 2-S3 A) Schematic mechanism of allopurinol and function of XDH protein in cells

B) A theoretical toy model (A Fuzzy Metabolic Switch) that can explain addiction to XDH protein in allopurinol-sensitive cells which have higher levels of basal XDH protein. Based on the genomic signatures and enrichment analysis these cell lines can be more dependent to Pentose Phosphate Pathway (PPP) by increasing their XDH protein level while resistant cell lines can be more dependent on Fatty Acid Oxidation (FAO) and Glycolysis. Allopurinol inhibits XDH protein leading to a metabolic stress and cell death.

Figure 2-S4 Western blot showing the protein levels of JAK2 s after siRNA knockdown. Figure 2-S5 CI for different doses of CEP-33779 and allopurinol for 3 resistant cell lines. Figure 2-S6 A and B) Cell viability after treatment with allopurinol, CEP-33779 and combined allopurinol and CEP-33779 represented by proliferation in soft agar gel in one resistant and one sensitive cell line (Allopurinol 400μM and CEP-33779 1.6μM). The quantification in panel B is presented as mean±SEM. Comparison was done between all 3 treatments and vehicle-control (**p<0.01, ****p<0.0001)

Figure 2-S7 Changes in the body weight of mice used as xenograft models for 3 different cell lines to evaluate combination therapy with CEP-33779 and allopurinol (mean±SEM)

Figure 3-S1: CI for different doses of allopurinol and CEP-33779 in NCI-H2170.

Figure 3-S2: Effects of combination treatment with CEP-33779 and allopurinol on cell viability of NCI-H2106. This cell line was inactive in response to single treatments and combination treatments.

Figure 4-S1: Images of 3 tumors of different treatment groups after treatment of 4 different PDX models. (Allopurinol (70 mg/kg daily), CEP-3379 (10 mg/kg daily), combination therapy (Allopurinol 50 mg/kg and CEP-33779 2.5 mg/kg daily) and PBS as placebo daily)

Figure 4-S2: Western blots of tumor samples after treatment course; apoptosis was measured by presence of cleaved caspase 3 in TM01563 and TM00188 models. (Allopurinol (70 mg/kg daily), CEP-3379 (10 mg/kg daily), combination therapy (Allopurinol 50 mg/kg and CEP-33779 2.5 mg/kg daily) and PBS as placebo daily)

Figure 4-S3: *In vivo* imaging in mice bearing PDX tumor models to detect apoptosis using a pancaspase assay done after the treatment course

A) TM01563 PDX Model

1: Mouse with no tumor and with no pan-caspase assay 2: Mouse with no tumor but receiving pan-caspase assay 3: Mouse from placebo treatment group receiving pan-caspase assay 4: Mouse from allopurinol treatment group receiving pan-caspase assay

B) TM00188 PDX Model

1 (I, II, III): Mouse with no tumor and with no pan-caspase assay

2 (I, II, III): Mouse with tumor and pan-caspase assay but not getting any treatment

3 (I, II, III): Mouse from placebo treatment group receiving pan-caspase assay

4 (I): Mouse from allopurinol treatment group receiving pan-caspase assay

4(II): Mouse from CEP-33779 treatment group receiving pan-caspase assay

4(III): Mouse from combination therapy group receiving pan-caspase assay

C) TM00206 PDX Model

1: Mouse with no tumor and with no pan-caspase assay 2: Mouse with tumor and pan-caspase assay but receiving no treatment 3: Mouse from placebo treatment group receiving pan-caspase assay 4: Mouse from allopurinol treatment group receiving pan-caspase assay

Figure 4-S4: Changes in body weights for the mice used for *in vivo* study of PDX tumor models

